# Evaluation of Retinal Safety of Hypoxia-Inducible Factor Prolyl Hydroxylase Inhibitors

**DOI:** 10.64898/2026.06.29.735161

**Authors:** Junki Hoshino, Kazuki Irie, Akimitsu Konishi, Hideo Akiyama, Yoji Andrew Minamishima

## Abstract

Hypoxia-inducible factor prolyl hydroxylase (HIF-PH) inhibitors are widely used for the treatment of renal anemia; however, their effects on intraocular vascular endothelial growth factor (VEGF) expression remain unclear. In this study, we examined the effects of all five HIF-PH inhibitors —roxadustat, daprodustat, vadadustat, enarodustat, and molidustat—on *Vegfa* expression in the retina in mice. C57BL/6J mice were orally administered each inhibitor. Six hours after administration, the kidney, retina, and liver were collected, and transcription levels were quantified by real-time quantitative reverse transcription PCR. Renal *Epo* transcription was significantly increased by molidustat (P < 0.01), roxadustat (P < 0.01), and enarodustat (P < 0.05). Retinal *Vegfa* transcription was significantly increased by four inhibitors (P < 0.01), with molidustat showing no significant effect. In the liver, *Vegfa* transcription was increased by daprodustat (P < 0.05) and vadadustat (P < 0.01). Furthermore, renal *Epo* and retinal *Vegfa* transcription levels showed a moderate positive correlation with a marginal trend toward statistical significance (r = 0.37, P = 0.08). These findings indicate that HIF-PH inhibitors differentially regulate hypoxia-responsive genes across tissues and suggest that retinal VEGF upregulation should be considered when evaluating the safety of these agents.

## Introduction

Chronic kidney disease (CKD) affects an estimated 800 million individuals worldwide and is frequently accompanied by renal anemia^1^, one of the major complications of the disease. Hypoxia-inducible factor-2 (HIF-2) plays a central role in erythropoiesis by regulating erythropoietin (EPO) production in response to hypoxia, primarily in the kidney and liver, while also regulating genes involved in iron uptake, storage, and transport to ensure adequate iron availability for red blood cell synthesis^2^. In CKD, impaired renal function leads to reduced EPO production, and chronic inflammation associated with uremia elevates hepcidin levels, thereby limiting iron utilization and aggravating anemia^3^. The current first-line therapy for renal anemia consists of erythropoiesis-stimulating agents (ESAs), but ESA therapy has been associated with increased cardiovascular risk and variable responsiveness^4^. In recent years, HIF-PH inhibitors have been developed as a novel therapeutic approach that stabilizes HIF under normoxic conditions, thereby enhancing endogenous EPO synthesis and improving iron metabolism, offering a more physiologically regulated alternative to conventional ESA therapy^2^.

Despite the treatment efficacy of HIF-PH inhibitors for anemia, chronic activation of HIF raises safety concerns. For instance, HIF-PH knockout mice develop severe cardiac hypertrophy resembling dilated cardiomyopathy, leading to premature death^5,6^. A major concern is the overproduction of vascular endothelial growth factor (VEGF), a glycoprotein involved in angiogenesis and vascular permeability^7^. Elevated VEGF expression is well documented in malignancies, where it correlates with tumor progression and poor prognosis^7^. Although concerns have been raised regarding the potential tumor-promoting effects of HIF activation, no clinical trials have demonstrated an increased risk of tumorigenesis associated with HIF-PH inhibitor treatment^8^. In the eye, VEGF also plays a pivotal role in the pathogenesis of chorioretinal diseases, including ischemia-related conditions such as diabetic retinopathy^9^ and retinal vein occlusion^10^, as well as neovascular age-related macular degeneration (nAMD)^11^. These conditions exhibit significantly elevated intraocular VEGF levels, which are closely associated with disease severity and progression. Although HIF-PH inhibitors may cause only mild increases in systemic VEGF levels^12^, their impact on intraocular VEGF expression and ocular angiogenesis remains largely unknown. To address this gap, this study aims to evaluate the effects of all five HIF-PH inhibitors currently approved for clinical use in Japan on retinal VEGF expression in mice. This investigation seeks to clarify the potential ocular risks associated with HIF-PH inhibitors and to inform the safe clinical use of these agents.

## Results

### Effects of HIF-PH Inhibitors on Kidney *Epo* Transcription

To investigate the pharmacological effects of HIF-PH inhibitors in mice, we first examined *Epo* expression levels in the kidney following drug administration. Each HIF-PH inhibitor was orally administered at a dose equivalent to the human maximum clinical dose, calculated using human equivalent dose (HED) conversions for mice (roxadustat; 15 mg/kg body weight, daprodustat; 5 mg/kg body weight, vadadustat; 120 mg/kg body weight, enarodustat; 1.5 mg/kg body weight, molidustat; 40 mg/kg body weight). For compounds that failed to induce *Epo* transcription at the initial dose, the dose was increased up to 3× the original dose. Vehicle control mice received 1% methyl cellulose orally. Six hours after administration, the mice were euthanized, and the kidneys were collected for analysis by real-time RT-PCR (Fig. 1A).

**Figure 1.**
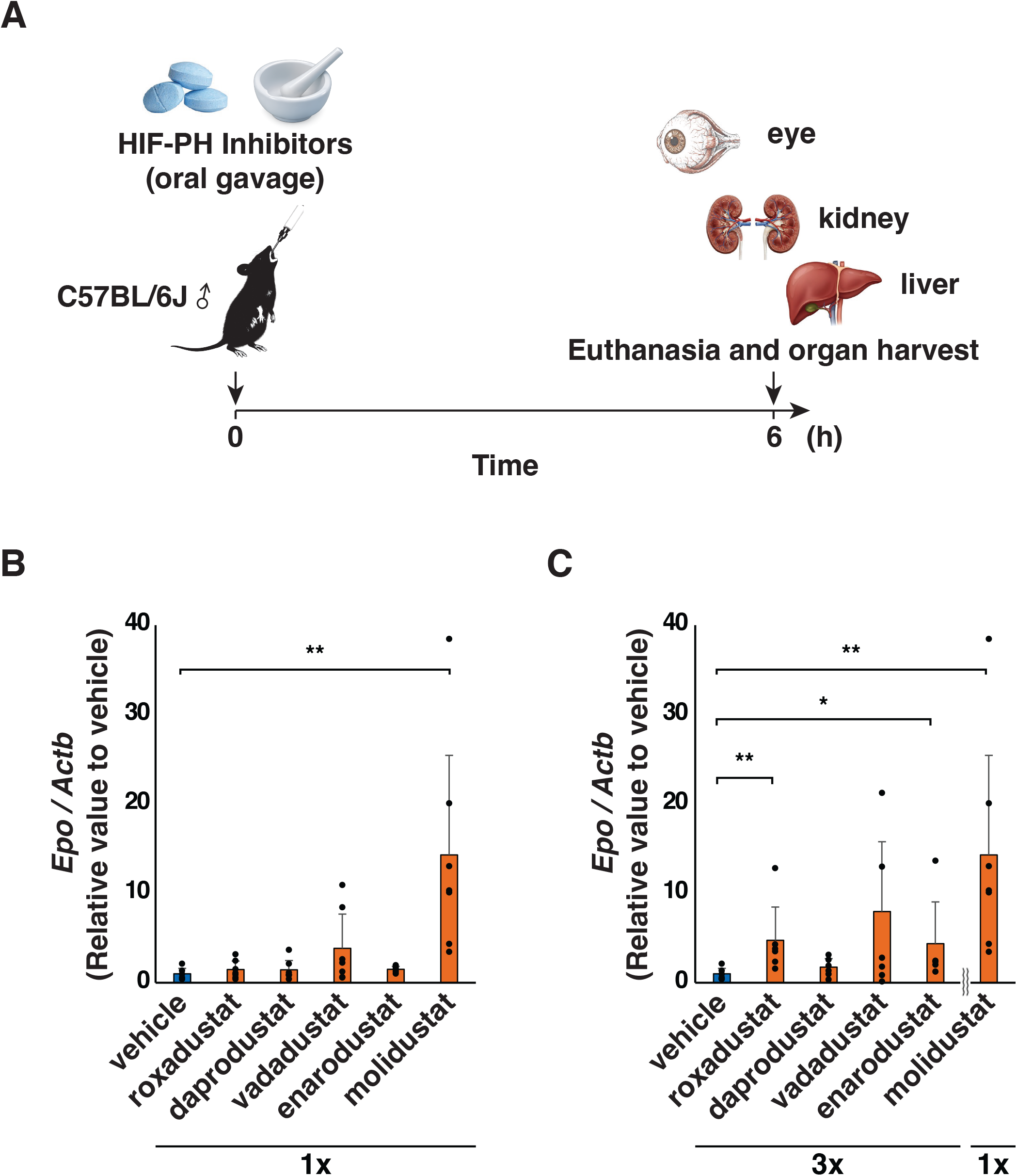
**A.** Schematic overview of the experimental timeline in mice. **B**. Renal *Epo* transcription following administration at the clinical-equivalent dose. A significant increase in *Epo* expression compared with the control was observed only with molidustat (** P < 0.01). **C**. Renal *Epo* transcription following clinical-equivalent dosing of molidustat and 3× dosing of other inhibitors. Significant increases in *Epo* expression compared with the control were observed with roxadustat (**P < 0.01), vadadustat (*P < 0.05), and molidustat (**P < 0.01).

At the 1× maximum clinical dose, only molidustat significantly increased *Epo* transcription in the kidney compared to control mice (P < 0.01) (Fig 1B). When the other four inhibitors were administered at 3× the dose, significant increases in *Epo* transcription were observed with roxadustat (P < 0.01) and enarodustat (P< 0.05), as well as with 1× molidustat. In contrast, daprodustat and vadadustat did not induce significant changes in *Epo* transcription at either dose level (Fig 1C).

### Effects of HIF-PH Inhibitors on *Vegfa* Transcription in the Retina and Liver

In the same group of mice used to assess renal *Epo* transcription, the retina and liver were also harvested to evaluate *Vegfa* transcription by real-time RT-PCR. To validate our experimental system for detecting hypoxia-induced gene expression, retinal explants from untreated mice were cultured *ex vivo* under normoxic or hypoxic conditions in a pilot study. As expected, *Vegfa* transcription was significantly increased under hypoxia compared with normoxia (P < 0.05; Fig. 2A), confirming the assay’s sensitivity to hypoxic stimulation.

**Figure 2.**
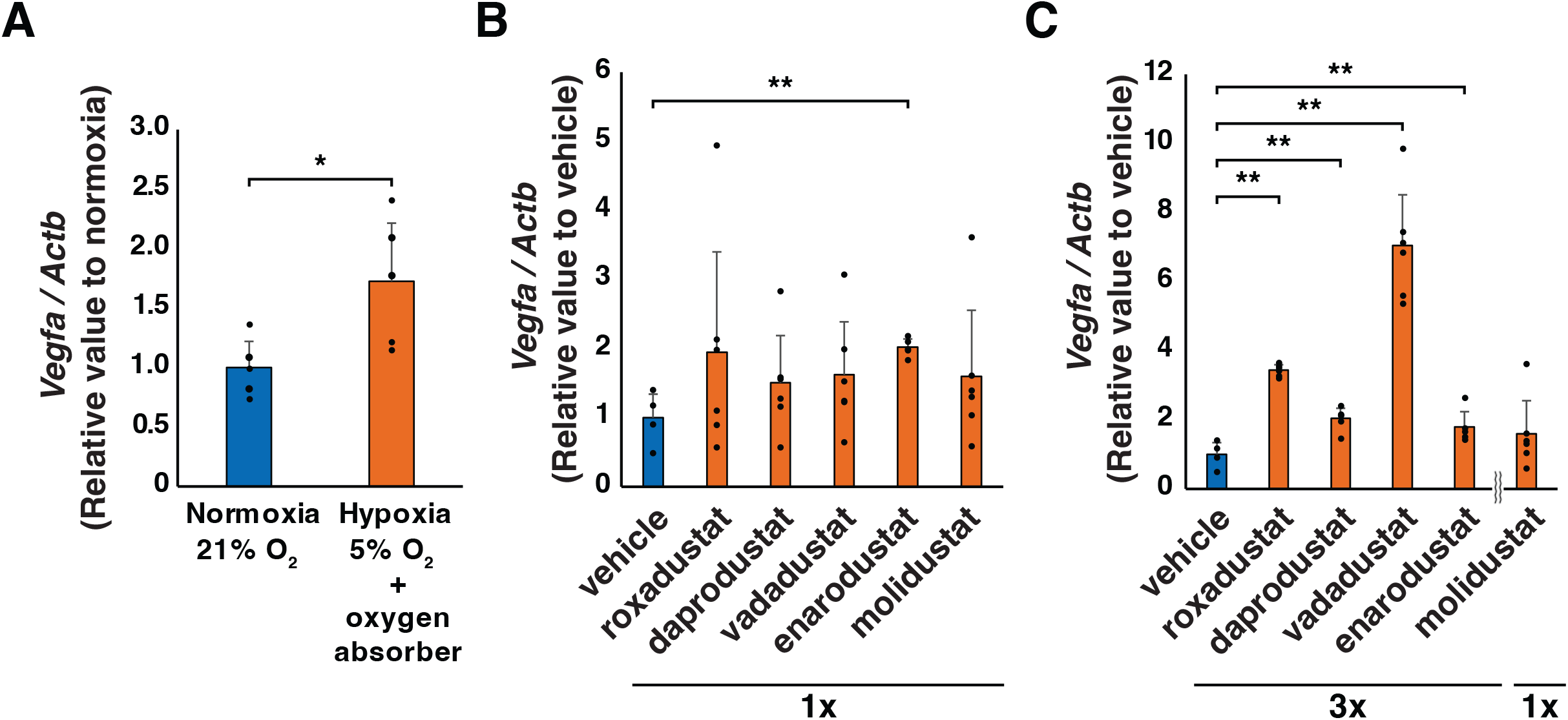
**A.** Comparison of *Vegfa* transcription in retinal explant cultured under hypoxic and normoxic conditions. *Vegfa* expression was significantly increased in retinas cultured under hypoxic conditions compared with those cultured under normoxic conditions (*P < 0.05). **B**. Retinal *Vegfa* transcription following administration at the clinical-equivalent dose. Only enarodustat significantly increased *Vegfa* expression compared with the control (P < 0.01). **C**. Retinal *Vegfa* transcription following clinical-equivalent dosing of molidustat and 3× dosing of other inhibitors. Compared with the control, *Vegfa* expression was increased by four of the five HIF-PH inhibitors (P < 0.01), excluding molidustat.

At the 1× maximum clinical dose, a significant increase in retinal *Vegfa* transcription was detected only with enarodustat (P < 0.01). (Fig 2B). At the 3× dose level, all four inhibitors other than molidustat significantly increased retinal *Vegfa* transcription (all P < 0.01) (Fig. 2B).

In the liver, none of the five HIF-PH inhibitors induced a significant change in *Vegfa* transcription at the 1× maximum clinical dose (Fig 3A). However, at the 3× dose level, daprodustat (P < 0.05) and vadadustat (P < 0.01) significantly increased hepatic *Vegfa* transcription compared to controls, while the other agents had no significant effect (Fig 3B).

**Figure 3.**
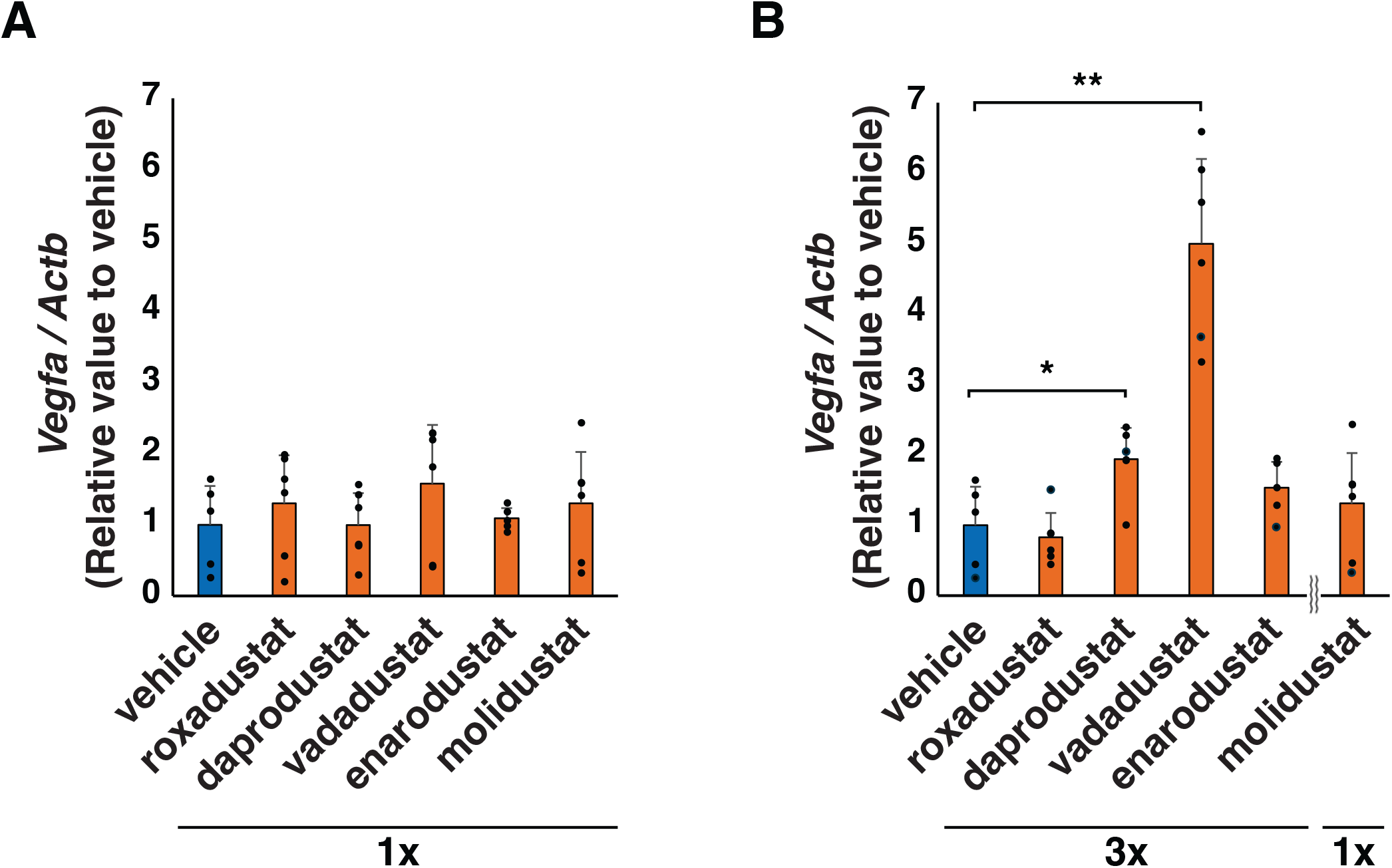
**A.** Hepatic *Vegfa* transcription following administration at the clinical-equivalent dose. No drug significantly increased *Vegfa* expression compared with the control. **B**. Hepatic *Vegfa* transcription following clinical-equivalent dosing of molidustat and threefold dosing of other inhibitors. Daprodustat (P < 0.05) and vadadustat (P < 0.01) significantly increased *Vegfa* expression compared with the control.

### Correlation Between Kidney *Epo* transcription and Retinal *Vegfa* transcription

The correlation between kidney *Epo* transcription and retinal *Vegfa* transcription was evaluated in mice treated with HIF-PH inhibitors at 3× the maximum clinical dose. In molidustat-treated mice, kidney *Epo* transcription showed high variability, and no significant increase in retinal *Vegfa* transcription was observed; therefore, this group was not included in the correlation analysis of the other treatment groups. Consequently, the correlation was assessed using data from the remaining four treatment groups, revealing a moderate positive correlation with a marginal trend toward statistical significance (r = 0.374, P = 0.079) (Fig. 4A). Because of the large variability observed in molidustat-treated mice, the correlation was analyzed separately for this group, and notably, despite the absence of a significant increase in retinal *Vegfa* transcription, a strong positive correlation trend between kidney *Epo* and retinal *Vegfa* transcription was also observed in molidustat-treated mice (r = 0.829, P = 0.058) (Fig. 4B).

**Figure 4.**
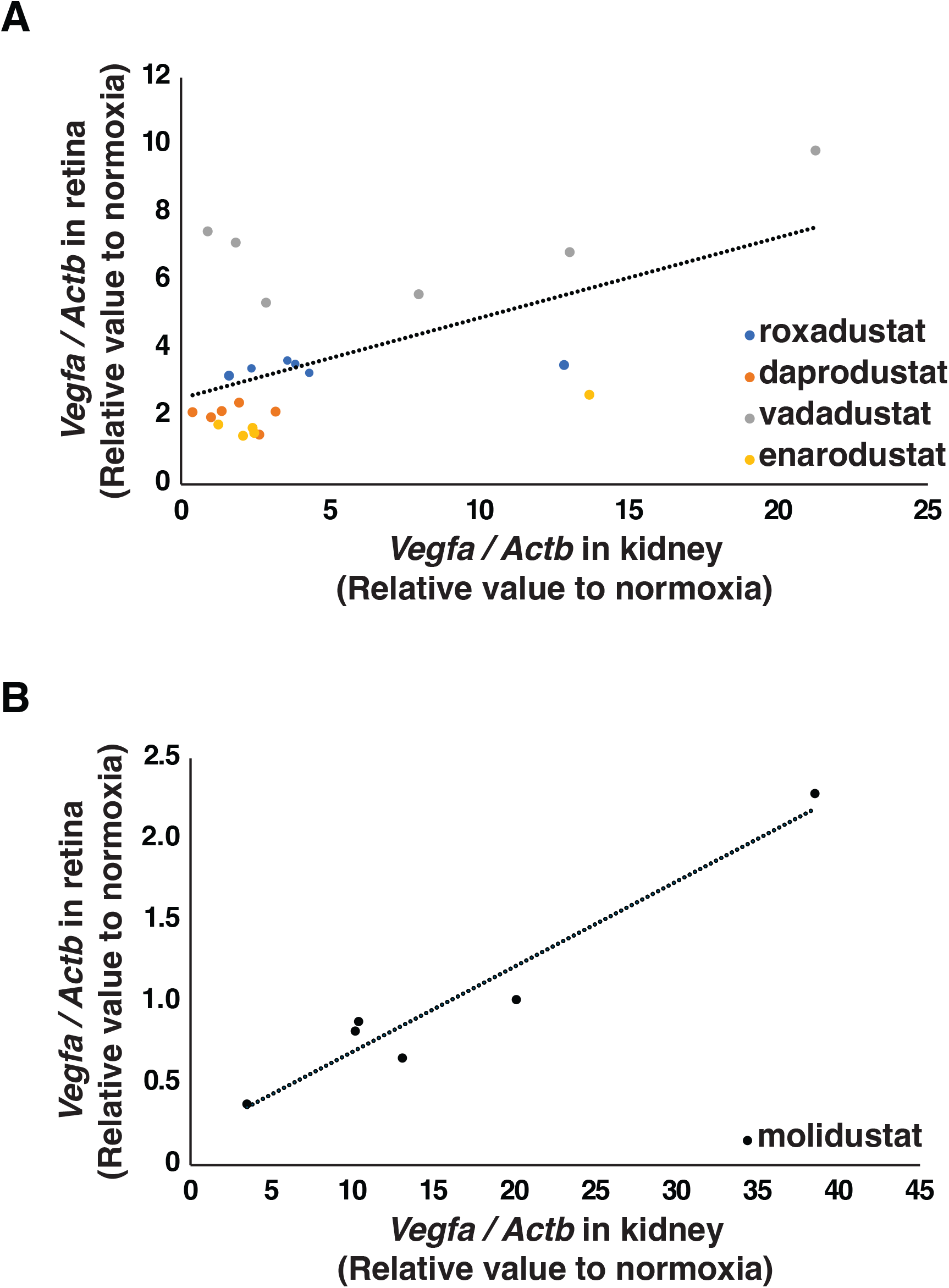
**A.** Correlation between renal *Epo* and retinal *Vegfa* transcription excluding molidustat. A moderate positive correlation trend was noted between renal *Epo* and retinal *Vegfa* transcription (r = 0.37, P = 0.08). **B**. Correlation between renal *Epo* and retinal *Vegfa* transcription in molidustat-treated mice (r = 0.829, P = 0.058). Dashed lines represent the regression lines.

## Discussion

In this study, we investigated the effects of HIF-PH inhibitors on retinal VEGF expression in mice. Six hours after administration, a significant upregulation of retinal *Vegfa* transcription was observed with four of the five agents, excluding molidustat. Furthermore, the extent of this upregulation tended to correlate positively with the increase in *Epo* transcription in the kidney.

Diabetic nephropathy is one of the leading causes of CKD and one of the major complications of diabetes. Among the other major diabetic complications is diabetic retinopathy. The development of diabetic retinopathy is multifactorial, driven by factors such as retinal ischemia and chronic inflammation, with treatment primarily aimed at suppressing VEGF production^9^. Intraocular VEGF levels are known to be higher in eyes with diabetic retinopathy than in those without, and these levels increase with disease progression^13^. For advanced stages of the disease, panretinal photocoagulation is performed to reduce VEGF production, and intravitreal injection of anti-VEGF agents is the standard treatment for diabetic macular edema^14^. Although large clinical trials of HIF-PH inhibitors have not reported adverse events such as the onset or worsening of diabetic retinopathy or diabetic macular edema, several case reports have described such associations^15,16^. In light of these reports and the findings of the present study, caution may be warranted when administering HIF-PH inhibitors to patients with renal anemia and coexisting diabetic retinopathy. Furthermore, although no clinical cases have been reported to date, the results of this study—demonstrating increased retinal VEGF expression following HIF-PH inhibitor administration— raise the possibility that disease progression could also be exacerbated in patients with other VEGF-dependent ocular diseases, such as retinal vein occlusion and neovascular age-related macular degeneration.

In the present study, vadadustat induced the greatest upregulation of retinal *Vegfa* transcription among the five HIF-PH inhibitors evaluated. Vadadustat has been investigated in two phase 3 clinical trials: the INNO2VATE trials, which targeted patients with dialysis-dependent chronic kidney disease^17^, and the PRO2TECT trials, which enrolled patients with non-dialysis-dependent CKD^18^. Both trials assessed the efficacy and safety of vadadustat in comparison with the ESA darbepoetin alfa for the treatment of renal anemia. In both trials, vadadustat demonstrated non-inferiority to darbepoetin alfa with respect to erythropoietic efficacy. Regarding cardiovascular safety, as measured by the incidence of major adverse cardiovascular events (MACE), vadadustat met the predefined non-inferiority margin in the INNO2VATE^17^, but failed to do so in the PRO2TECT trials^18^. Among the five HIF-PH inhibitors examined in this study, vadadustat was the only agent for which non-inferiority to ESA in terms of MACE risk was not demonstrated in a large-scale clinical trial^18-26^. The marked upregulation of retinal *Vegfa* transcription observed with vadadustat in this study suggests that hypoxia-responsive pathways mediated by HIF activation may be more broadly activated in tissues beyond the intended target organs such as the kidney and liver. This off-target HIF activation could potentially contribute to both the increased retinal *Vegfa* transcription observed in this study and the elevated cardiovascular risk reported in clinical trials.

In this study, administration of molidustat significantly increased renal *Epo* transcription, whereas no notable changes were observed in retinal *Vegfa* transcription. Molidustat is unique among HIF-PH inhibitors in that it does not possess a 2-oxoglutarate (2-OG) backbone, a structural motif shared by most other inhibitors and typically involved in competitive binding to the catalytic site of HIF-PH (PHD) enzymes^27^. Owing to this structural difference, molidustat is thought to inhibit HIF-PH through a mechanism that is partially distinct from that of other agents in this class. Such mechanistic divergence may influence the pattern and magnitude of HIF stabilization and downstream gene activation. In this context, it is conceivable that molidustat induces a more modest activation of pathways related to VEGF regulation, resulting in the relatively limited upregulation of retinal *Vegfa* transcription observed in our study. These findings suggest that structural and mechanistic heterogeneity among HIF-PH inhibitors may translate into differences in ocular biological effects, underscoring the need to consider drug-specific properties when evaluating their potential ocular safety.

A major feature of this study is that the dose of each agent was set according to the clinical dose used in humans. As reported by Schofield et al., vadadustat showed markedly lower potency in cellular assay systems compared with other HIF-PH inhibitors, with differences in cellular uptake or metabolism suggested as possible contributing factors^27^. Although *in vivo* factors are also likely to play a role in this discrepancy, their findings are consistent with the clinical dose at which vadadustat is administered at higher levels than other agents. Taken together, these results indicate that adjusting dosing regimens to achieve equivalent pharmacological effects across different agents is not straightforward^28^, making direct comparisons challenging. A major strength of this study is the use of clinically based dosing as the foundation of the experimental design, particularly with respect to evaluating safety in clinical settings.

In addition to differences in HIF-PH inhibitory potency and pharmacokinetic profiles, off-target effects unrelated to canonical HIF stabilization may also contribute to the distinct biological responses observed among HIF-PH inhibitors. Roxadustat, for example, has been reported to share structural similarity with triiodothyronine (T3), and clinical studies have demonstrated reductions in serum thyroid-stimulating hormone (TSH) and free thyroxine (free T4) levels following its administration. These findings raise the possibility that certain HIF-PH inhibitors may interact with molecular pathways beyond the hypoxia response system^29,30^.

Furthermore, most HIF-PH inhibitors are structurally related to 2-oxoglutarate (2-OG) and act as competitive inhibitors of 2-OG-dependent dioxygenases. Although these agents are designed to selectively inhibit HIF-PH (PHD) enzymes, the human genome encodes many 2-OG-dependent enzymes, including Jumonji C-domain-containing histone demethylases, lysine demethylases (KDMs), ten-eleven translocation (TET) DNA demethylases, and other iron- and 2-OG-dependent oxygenases. Depending on drug concentration, tissue distribution, and the relative affinity of individual enzymes for 2-OG and each inhibitor, unintended modulation of these enzymes may occur. Such effects could influence transcriptional regulation, epigenetic states, and cellular metabolism independently of canonical HIF signaling^31^.

In this context, the differential retinal *Vegfa* transcription responses observed among the five HIF-PH inhibitors in the present study may not be explained solely by differences in HIF stabilization and could also reflect broader pharmacological heterogeneity among these compounds. Further studies will be necessary to determine whether non-canonical targets contribute to the ocular and systemic effects of HIF-PH inhibitors.

This study has several limitations. First, the experiments were conducted in mice, and the systemic exposure and tissue distribution of HIF-PH inhibitors may differ from those in humans. Therefore, the retinal responses observed in this study may not fully reflect the clinical situation. Second, gene expression was assessed only at a single time point, and dynamic changes in *Vegfa* transcription following drug administration were not evaluated. Third, we did not measure protein levels of EPO and VEGF, or perform functional assessments, such as vascular permeability or retinal imaging, to directly determine the biological impact of HIF activation. Finally, the doses of each inhibitor were based on clinical equivalents; however, differences in pharmacokinetics between individual agents may influence the comparability of the findings.

In conclusion, the present study indicates that HIF-PH inhibitors have the potential to increase retinal VEGF expression. These results highlight the need for caution when using these agents in individuals with VEGF-dependent ocular diseases.

## Materials and Methods

### Mice

All experiments were approved by the Animal Care and Utilization Committee of the Gunma University School of Medicine (23-034). Wild-type mice used in this study were C57BL/6J male mice aged 8 weeks, which were obtained from CLEA Japan, Inc. The mice were maintained on a 12:12-h light-dark cycle in a specific-pathogen-free facility and were fed ad libitum.

### Treatment of mice using HIF-PH inhibitors

Mice were treated with either the vehicle (1% methyl cellulose 400) (132-05055; FUJIFILM Wako, Osaka, Japan) or HIF-PH inhibitors: roxadustat (Astellas, Tokyo, Japan), daprodustat (GlaxoSmithKline, London, UK), vadadustat (Mitsubishi Tanabe, Osaka, Japan), enarodustat (JAPAN TOBACCO, Tokyo, Japan), or molidustat (Bayer, Leverkusen, Germany) by oral gavage. Each commercially available tablet was milled and suspended in 1% methyl cellulose 400 for oral administration.

### Retinal explant culture

Untreated mice were euthanized, and the retinas were dissected from both eyes. One retina from each mouse was incubated under normoxic conditions (21% O_2_), while the fellow retina was incubated under hypoxic conditions (5% O_2_) in a sealed container containing an oxygen absorber. Both retinas were cultured for 6 hours at 37°C in high-glucose medium (DMEM 043-30085; FUJIFILM Wako, Osaka, Japan) supplemented with 10% fetal bovine serum (SIGMA #173012).

### Total RNA extraction and real-time RT-PCR analysis

Total RNA was extracted from the retinas, livers, and kidneys of each mouse using ReliaPrep RNA Tissue Miniprep Systems (Z6011; Promega, Madison, US). Reverse transcription and Real-time PCR (real-time RT-PCR) were performed using ReverTra Ace qPCR RT Master Mix (FSQ-201; Toyobo, Osaka, Japan) and Thunderbird Next SYBR qPCR mix (QPX-201; Toyobo, Osaka, Japan) with StepOnePlus Real-Time PCR System (Thermo Fisher Scientific, Massachusetts, US). The mRNA levels were normalized to the *Actb* level. The primer sequences^32-34^ are shown in Table 1.

**Table 1.**
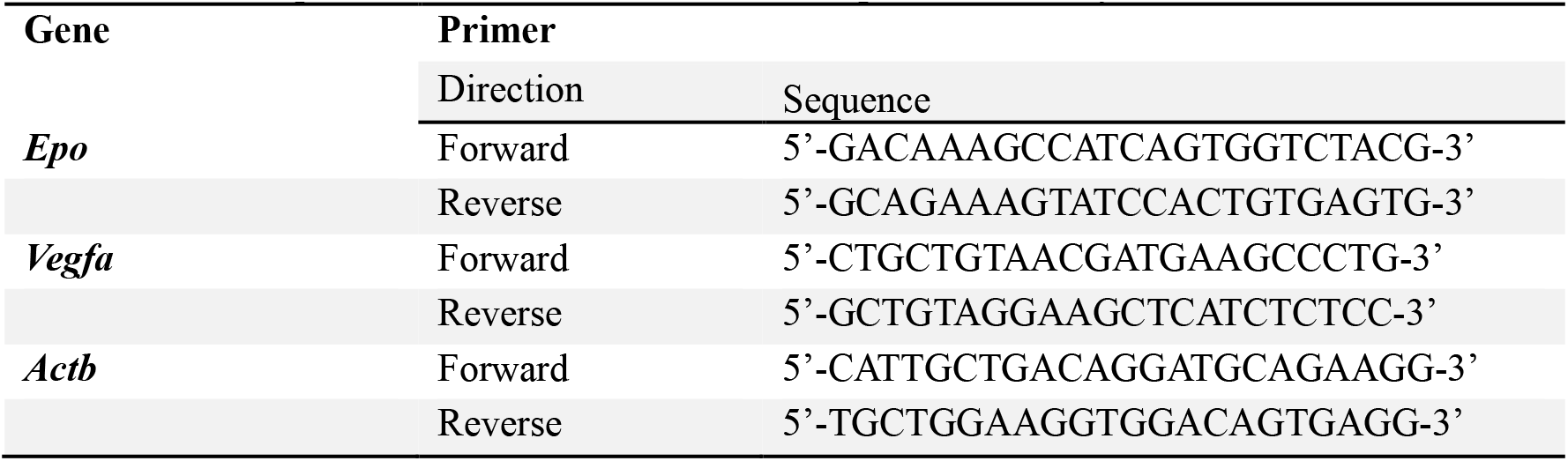
Primer sequences for real-time reverse transcription-PCR analysis.

## Statistics

Multiple comparisons between control mice and each drug-treated group were performed using Dunnett’s test. Comparisons between retinas cultured under normoxic and hypoxic conditions were analyzed using the Mann-Whitney U test. The correlation between kidney *Epo* mRNA and retinal *Vegfa* transcription levels was assessed using Spearman’s rank correlation coefficient. P < 0.05 was considered statistically significant. All statistical analyses were performed using R version 4.3.1 with RStudio version 2024.4.2.0.

## Acknowledgement

This work was performed using research equipment shared through the MEXT Project for Promoting Public Utilization of Advanced Research Infrastructure (Program for Supporting the Introduction of the New Sharing System, Grant Number JPMXS0420600124). The authors thank Chizuko Tomaru for administrative support, budget management, and assistance with manuscript preparation.

## Author Contributions

J.H., K.I., A.K., and Y.A.M. designed the research. J.H. conducted the research. J.H., H.A., and Y.A.M. wrote the paper.

## Data availability statement

The datasets used and/or analyzed during the current study are available from the corresponding author upon reasonable request.

